# Heterogeneous nanoplastic exposure induces integrated immunometabolic states in human monocytes under physiological flow

**DOI:** 10.64898/2026.03.11.711031

**Authors:** Vladimir Kovacevic, Nevena Milivojević Dimitrijević, Marija Mihailovich, Marko Zivanovic, Milos Ivanovic, Andreja Zivic, Marina Gazdic Jankovic, Aleksandra Kovacevic, Uršula Prosenc Zmrzljak, Feđa Puac, Nenad Filipović, Biljana Ljujic

## Abstract

Nanoplastics interact continuously with circulating immune cells, yet how particle size and exposure complexity shape immune transcriptional organization under physiological flow conditions remains poorly understood. Here, controlled microfluidic exposure was combined with single-cell RNA sequencing to investigate the effects of size-defined polystyrene nanoplastics (PSNPs; 40 nm, 200 nm, and 40 + 200 nm) on primary human peripheral blood mononuclear cells (PBMCs) under dynamic flow conditions. Across immune populations, PSNP exposure induced a conserved ribosome-associated and RNA-regulatory transcriptional program, indicating a shared intracellular adaptive response. Monocytes displayed the strongest transcriptional remodeling, characterized by coordinated modulation of ribosome-associated, metabolic, and inflammatory signaling pathways in a size-dependent manner. Exposure to 40 nm PSNPs negatively enriched (suppressed) mitochondrial metabolic pathways, whereas 200 nm PSNPs enriched inflammatory signaling programs. Combined exposure induced concurrent metabolic and inflammatory pathway engagement without evidence of major immune topology disruption or discrete inflammatory state transitions. In contrast, adaptive immune cells exhibited comparatively modest and lineage-preserving transcriptional modulation. Together, these findings demonstrate that nanoplastic size and exposure complexity shape coordinated immunometabolic adaptation in human immune cells under physiologically relevant flow conditions and establish a framework for studying dynamic material–immune interactions at single-cell resolution.

## 1. INTRODUCTION

Plastics have become an essential component of modern life due to their versatility, durability, and low cost. However, the environmental consequences of plastic pollution have gained substantial attention over recent decades, coinciding with a significant increase in the global health burden associated with environmental contaminants (Prata, 2018). Microplastics were identified in human blood for the first time in 2022, with 17 out of 22 individuals testing positive during screening (Kuhlman, 2022). Since then, increasing evidence has confirmed the presence of micro- and nanoplastics across a wide range of biological compartments (Sone et al., 2026). In parallel, the rapid rise in immune-mediated inflammatory diseases, particularly during transitions from developing to industrialized societies, highlights how environmental and lifestyle changes contribute to shifts in disease epidemiology (Toussaint et al., 2019).

Although environmental exposure to micro- and nanoplastics is generally considered chronic, several real-world scenarios can generate acute, high-particle-load exposures. Individuals may ingest up to 121,000 microplastic particles per year, with an additional ∼90,000 particles annually from bottled water consumption, while inhalation exposure may transiently reach ∼700,000 airborne fibers per washing machine cycle (Bojic et al., 2020). In addition, consumer products such as plastic tea bags and disposable coffee cups can release up to 3.1 × 10⁹ nanoplastic particles and 1.26 × 10⁸ particles mL⁻¹, respectively (Hernandez et al., 2019; Lambert & Wagner, 2016). Microwave heating of plastic containers has further been shown to release up to 2.11 × 10⁹ nanoplastic particles and 4.22 × 10⁶ microplastic particles per cm² within minutes (Hussain et al., 2023). Yet, the biological consequences of acute, high-dose exposure to micro- and nanoplastics remain poorly understood.

Micro- and nanoplastics can enter the human body through inhalation, ingestion, and potentially dermal exposure, where they may interact with cells and tissues at the molecular level (Prata, 2018; Toussaint et al., 2019; Wu et al., 2022). Toxicokinetic studies suggest that, once internalized, nanoplastics can be systemically distributed and persist in cells and organs, raising concerns about long-term biological effect (Wolff et al., 2023; Wu et al., 2022). Micro- and nanoplastics have been detected in multiple human tissues, underscoring their capacity for systemic exposure and cellular interaction (Marcelino et al., 2022; Nihart et al., 2025; Ragusa et al., 2022; Ragusa et al., 2021; Zheng et al., 2024). Experimental evidence increasingly indicates that nanoplastics can induce oxidative stress, perturb cellular metabolism, and modulate inflammatory pathways (Antunes et al., 2023; Mahmud et al., 2024; Yee et al., 2021; Zhang et al., 2022). These responses differ across cell types and are influenced by particle size, surface chemistry, and exposure context.

Our group previously demonstrated that polystyrene nanoparticles (PSNPs) can alter transcriptional programs in human developmental models and, upon oral administration in mice, induce systemic effects, including neurobehavioral alterations and immune activation (Bojic et al., 2020; Nikolic et al., 2022). *In vivo* exposure to mixed-size PSNP populations increased pro-inflammatory IL-12- and IL-23-producing CD3+ cells and reduced anti-inflammatory IL-10-producing CD3+ cells in the spleen, supporting the immunomodulatory potential of PSNPs. Several studies further demonstrated that nanoplastics can be internalized by immune cells, including macrophages and B lymphocytes, leading to mitochondrial dysfunction, increased reactive oxygen species (ROS) production, and disruption of metabolic pathways (Rajendran & Chandrasekaran, 2023).

Wolff and colleagues, using primary human immune cells, demonstrated that polystyrene (PS) micro- and nanoplastics induce distinct effects across innate and adaptive immune populations in a size- and polymer-dependent manner (Wolff et al., 2023). T cells isolated from human peripheral blood mononuclear cells (PBMCs) were relatively resistant to cytotoxicity following PS particle exposure but exhibited increased expression of activation markers after 72 h, along with strongly altered cytokine secretion profiles, indicating functional immune modulation in the absence of pronounced cell death. In contrast, monocyte-derived macrophages and dendritic cells displayed substantially higher sensitivity to micro- and nanoplastics. Exposure of these innate immune cells for 24 h induced changes in inflammatory surface marker expression and suppression of inflammatory phenotypes, consistent with polarization toward an M2-like macrophage state. Together, these findings demonstrated that PS particles are sufficient to alter immune cell activation and inflammatory programs in a cell-type–specific manner.

Recent studies further demonstrated that PSNPs directly interact with primary human blood immune cells in a strongly cell-type- and size-dependent manner (Fusco et al., 2025). Using single-cell mass cytometry, Fusco et al. showed that PSNPs were preferentially associated with monocytes and dendritic cells, which also displayed the strongest reductions in viability and inflammatory activation following exposure. Larger particles (200 nm) induced more pronounced immune responses than 50 nm PSNPs, including increased CD25/CD69 expression and elevated production of pro-inflammatory cytokines such as TNF-α, IFN-γ, GM-CSF, and IL-4. In contrast, T cells exhibited comparatively limited cytotoxicity despite detectable functional modulation. Follow-up transcriptomic and proteomic analyses of THP-1 monocytic cells exposed to PSNPs for 24 h further linked PSNP exposure to oxidative stress, inflammatory signaling, and mitochondrial stress responses.

While microfluidic platforms have been widely used to study nanoparticle transport, separation, and detection, their application to investigate biological responses to nanoplastics remains extremely limited. Most studies examining the effects of PSNPs on human cells rely on static *in vitro* systems or bulk analyses, which do not recapitulate the dynamic conditions of circulation and provide limited insight into lineage-specific transcriptional organization and immune heterogeneity. Recent studies using primary human immune cells have demonstrated that PSNP responses depend on particle size and immune cell identity, with monocyte-lineage cells showing greater sensitivity than adaptive immune populations. However, how these parameters shape coordinated cell-type–specific transcriptional responses remains poorly defined, particularly under physiologically relevant flow conditions. In addition, it remains unresolved whether heterogeneous nanoplastic mixtures induce additive responses or whether immune cells integrate multiple material cues into unified transcriptional configurations. To our knowledge, the integration of controlled microfluidic exposure, primary human PBMCs, and single-cell transcriptomics to investigate immune responses to nanoplastics has not been previously reported.

Here, we combined controlled microfluidic exposure with single-cell RNA sequencing (scRNA-seq) to investigate how nanoplastic size and exposure complexity shape immune transcriptional responses in human PBMCs under physiological flow conditions. Using carboxylated PSNPs of defined sizes (40 nm and 200 nm) and their combination, we examined transcriptional organization across innate and adaptive immune populations at single-cell resolution. We identified cell-type–specific immune responses, with monocytes displaying the strongest transcriptional remodeling characterized by coordinated modulation of ribosome-associated and metabolic pathways. Our approach further enabled the systematic assessment of size-dependent responses, shared intracellular programs, and the extent to which mixed particle exposure generates additive versus integrated transcriptional states.

## 2. Materials and Methods

### 2.1 Ethics Statement

The study involving human-derived material was approved by the Ethics Committee of the University Clinical Center Kragujevac (UKC Kragujevac), Zmaj Jovina 30, 34000 Kragujevac, Serbia (approval number: 01/26-105). The study was conducted in accordance with the principles of the Declaration of Helsinki. Written informed consent was obtained from the donor prior to sample collection, and the signed consent forms are held at UKC Kragujevac.

### 2.2 Human peripheral blood mononuclear cell isolation

PBMCs were obtained from a healthy female donor (31 years old) after obtaining informed consent, in accordance with institutional and national ethical guidelines (Ethics Committee of the University Clinical Center Kragujevac, Serbia, approval number: 01/26-105). Whole blood was collected into EDTA-coated tubes and processed immediately. PBMCs were isolated by density gradient centrifugation with Ficoll-Paque PLUS (Cytiva) according to the manufacturer’s instructions. After centrifugation, the mononuclear cell layer was collected, washed with phosphate-buffered saline (PBS), and resuspended in RPMI-1640 medium (Gibco®) supplemented with 10% fetal bovine serum (FBS, Gibco®).

### 2.3 Microfluidic culture and nanoplastic exposure under flow

PBMCs were cultured on a microfluidic chip platform designed to enable immune cell culture under continuous-flow conditions. To support three-dimensional culture of non-adherent immune cells, PBMCs were embedded in hESC-qualified Matrigel® (Corning®, lot No 1332001) and introduced into the microfluidic channels at a density of approximately 10 × 10^6^ cells/ml. Cells were maintained under continuous flow throughout the experiment.

Cells were exposed for 24 h to carboxylated PSNPs under three experimental conditions: (i) 40 nm PSNPs, (ii) 200 nm PSNPs, or (iii) a mixed condition containing both 40 nm and 200 nm PSNPs. Final concentrations were 100 µg mL⁻¹ for single-size exposures and 50 µg mL⁻¹ each for mixed exposures. While these concentrations exceed estimated environmental exposure levels, they are consistent with prior mechanistic nanoplastic studies (Nacka-Aleksic et al., 2025; Nikolic et al., 2022), enabling characterization of early transcriptional adaptation under controlled conditions. Control samples were cultured under identical microfluidic conditions in the absence of PSNPs. Flow rate and shear stress were selected to approximate physiological conditions (0.03 ml/min).

### 2.4 Recovery of cells and viability assessment

After exposure, PBMCs were recovered from the Matrigel matrix using Cell Recovery Solution (Corning®) according to the manufacturer’s protocol. Recovered cells were filtered through a 40 µm cell strainer, washed, and resuspended in PBS. Cell counts and viability were assessed using trypan blue exclusion, with post-exposure viability exceeding 90% across all conditions.

### 2.5 Single-cell RNA sequencing

Single-cell RNA sequencing libraries were prepared using the Chromium Single Cell 3′ platform (10x Genomics). Approximately 10,000 cells were loaded per sample. Library preparation was performed according to the manufacturer’s instructions. Libraries were sequenced on an Illumina NovaSeq 6000 with a target sequencing depth of approximately 20,000 reads per cell and 138 sequencing cycles.

### 2.6 Preprocessing and quality control of single-cell RNA sequencing (scRNA-seq) data

Raw sequencing data were processed using Cell Ranger 7.0.1 for demultiplexing, alignment to the human reference genome (GRCh38), and generation of gene-cell count matrices. Downstream analyses were performed using Python-based single-cell analysis tools.

Cells with fewer than 200 genes were filtered to remove low-quality cells. Genes expressed in fewer than 12 cells were excluded. Data were normalized and log-transformed prior to downstream analyses.

### 2.7 Dimensionality reduction and cell type annotation

Uniform Manifold Approximation and Projection (UMAP) was used to visualize cellular heterogeneity. Projection of gene expression from all four samples into the UMAP plane (Supplementary Figure 1) revealed noticeable batch effects, as evidenced by Silhouette (Rousseeuw, 1987) and LISI (Korsunsky et al., 2019) scores of 0.207 and 1.08, respectively. For its removal, we used Harmony (Korsunsky et al., 2019), producing embeddings projected onto the UMAP plane without visible or measurable batch effects (Silhouette score of −0.001 and LISI score of 1.3).

Cell type annotation was performed using the reference-based Seurat (Butler et al., 2018) and CoDi-annotation tools (Vladimir Kovacevic, 2024). Major immune populations, including monocytes, B cells, CD4⁺ T cells, CD8⁺ T cells, and natural killer (NK) cells, were identified. The annotation strategy with the highest concordance was used for all downstream analyses. The Seurat tool (FindTransferAnchors method (Hao et al., 2021; Hao et al., 2024) was used to map cell types from the reference single-cell RNA dataset. Additionally, to validate the quality of the annotations, we used CoD (Vladimir Kovacevic, 2024) and its CoDi distance algorithm, both embedded in the same package.

Since the annotations differ, we decided to use literature-confirmed marker genes for the cell types obtained and check which annotations retain them as marker genes. For that purpose, we collected the following set from different sources:

- B cell (Liu et al., 2022): *BANK1, BACH2, CD19, CD22, CD37, CD74, CD79A, CD79B, CR2, FCER2, HLA-DRA, MS4A1, PAX5, SPIB, TNFRSF13B, TNFRSF13C*
- Monocyte (Hao et al., 2021): *CD14, LYZ, S100A8, S100A9, FCN1, CCR2, CSF3R, FCGR3A, ITGAM, CX3CR1, LST1, LGALS3, CTSS, TREM1, IL1B, IFI27, TNF, ICAM1, TLR4*
- CD4+ (Filippov et al., 2024; Liu et al., 2022): *CD3D, CD3E, CD3G, CD4, IL7R, CCR7, LTB, SELL, TCF7, LEF1*
- CD8+ (Grossman et al., 2004): *CD8A, CD8B, GZMA, GZMB, GZMK, PRF1, GNLY, NKG7, KLRD1, CTSW, CD2, CD27, IFNG*
- NK (Bongen et al., 2018; van Vliet et al., 2024): *NCAM1, NKG7, GNLY, KLRD1, KLRF1, GZMA, GZMB, GZMK, PRF1, FCGR3A, FGFBP2, KLRC1, NCR1, NCR3, XCL1, XCL2, TBX21, ZBTB16*

We annotated all four samples with these three tools and calculated marker genes for each cell type. Additionally, we identified marker genes for the reference scRNA used to validate our samples. Table 1 shows the percentages of captured literature-confirmed markers for each annotation. Since the CoDi_dist algorithm obtained the highest average capture rate, we decided to use its annotation for the downstream analysis.

**Table 1:** Scores (from 0 to 1) for capturing literature-validated marker genes for detected cell types. Seurat did not detect Natural killer cells, so the score is unavailable.

| annotation | B cell | Monocyte | CD4+ | CD8+ | NK | Average |
| --- | --- | --- | --- | --- | --- | --- |
| scRNA reference | 0.44 | 0.42 | 0.2 | 0.69 | 0.5 | 0.45 |
| CoDi | 0.62 | 0.1225 | 0.45 | 0.4025 | 0.36 | 0.391 |
| CoDi_dist | 0.62 | 0.1225 | 0.45 | 0.44 | 0.4275 | 0.412 |
| Seurat | 0.62 | 0.1225 | 0.45 | 0.42 | N/A | 0.4031 |

### 2.8 Differential gene expression analysis

Differential gene expression analyses were performed for each immune cell type, comparing PSNP-exposed cells to control cells. Statistical testing was performed using the Wilcoxon rank-sum test. P values were corrected for multiple testing using the Benjamini–Hochberg method, and genes with an adjusted p value < 0.05 were considered significantly differentially expressed. Although several studies have proposed the use of log-fold change thresholds (for example, logFC > 0.25) to score and prioritize differentially expressed genes (DEGs) (Satija et al., 2015), our analyses revealed substantial variability and instability of log-fold change values, particularly when comparing estimates derived from raw versus normalized expression data. To ensure consistency and reduce bias introduced by normalization procedures, DEGs in this study were therefore selected solely based on statistical significance (p-value).

### 2.9 Pathway enrichment analysis

Pathway enrichment analysis was performed using gene set enrichment analysis (GSEA) against KEGG pathway annotations from the GSEApy tool (Fang et al., 2023). Gene sets from the KEGG 2021 Human database (Kanehisa, 2019; Kanehisa et al., 2025; Kanehisa & Goto, 2000) were tested using 1,000 permutations to assess enrichment significance. Pathways with False discovery rate (FDR) q-values ≤ 0.05 were considered significant (Zhang et al., 2023). To ensure biological relevance and robustness, only gene sets with 10-500 contributing genes were retained. Each of these steps was implemented using Python libraries and tools commonly used in scRNA-seq data analysis, as detailed in the referenced Jupyter Notebook, including scanpy (Wolf et al., 2018) and GSEAP (Fang et al., 2023). This comprehensive approach ensures a robust and reproducible analysis of the impact of PSNPs on single-cell gene expression profiles. KEGG pathways were grouped into predefined functional modules, applied uniformly across cell types and exposure conditions.

### 2.10 Definition of biological modules

Enriched KEGG pathways were grouped into higher-order biological modules based on shared functional annotations and established biological relationships, enabling consistent interpretation across cell types and exposure conditions. Pathways related to ribosomal function, RNA processing, and transcriptional control were grouped under **Translation & RNA regulation**. Pathways associated with oxidative phosphorylation, the tricarboxylic acid (TCA) cycle, pentose phosphate pathway, fatty acid metabolism, and thermogenesis were grouped under **Mitochondrial metabolism and bioenergetics**. Vesicle trafficking and intracellular transport pathways were grouped under **Interface-associated cellular processes**. Canonical innate immune signaling pathways, including TNF and IL-17 signaling and virus–cytokine interaction pathways, were grouped under **Innate immune sensing & inflammatory signaling**. *The enrichment of the ‘Viral protein interaction with cytokine and cytokine receptor’ pathway reflects a shared transcriptional signature associated with sterile danger-associated molecular pattern (DAMP) sensing and intracellular stress-response cascades, rather than active viral processes.* Antigen processing and presentation pathways were grouped under **Antigen processing and immune communication**. Additional modules describing adaptive immune receptor signaling and cell adhesion, migration, and tissue interaction were defined for analyses of adaptive immune cell populations.

### 2.11 Trajectory and state transition analyses

To assess whether PSNP exposure induced continuous transcriptional trajectories or lineage transitions, diffusion pseudotime and partition-based graph abstraction (PAGA) analyses were performed using established single-cell trajectory inference methods from the scanpy (Wolf et al., 2018) framework. These analyses were used to evaluate global state transitions rather than to infer differentiation trajectories.

### 2.12 Data availability statement

Single-cell RNA sequencing data generated in this study are deposited in a public Zenodo repository (https://doi.org/10.5281/zenodo.15866724). The reference single-cell RNA dataset used for annotation was downloaded from the [Broad Institute’s repository](https://singlecell.broadinstitute.org/single_cell/study/SCP424/single-cell-comparison-pbmc-data#study-visualize).

### 2.13 Code availability

The source code for the analysis is publicly available on the GitHub repository at https://github.com/vladimirkovacevic/nano. The code for generating all the figures is available in notebooks/nano_figures.ipynb in the same repository.

## 3. RESULTS

### 3.1 24-hour exposure to PSNPs under physiological flow conditions does not alter immune composition

To investigate how PSNPs of different sizes and combinations modulate immune cell states under physiologically relevant conditions, we established a microfluidic exposure platform coupled to scRNA-seq (Figure 1). Peripheral blood–derived immune cells were isolated and exposed for 24 hours under continuous flow to monodisperse 40 nm PSNPs, 200 nm PSNPs, or a mixed exposure containing both particle sizes (40 + 200 nm), alongside unexposed controls. This platform enables controlled nanoparticle–cell interactions under dynamic conditions that better approximate physiological circulation than conventional static exposure paradigms.

**Figure 1.**
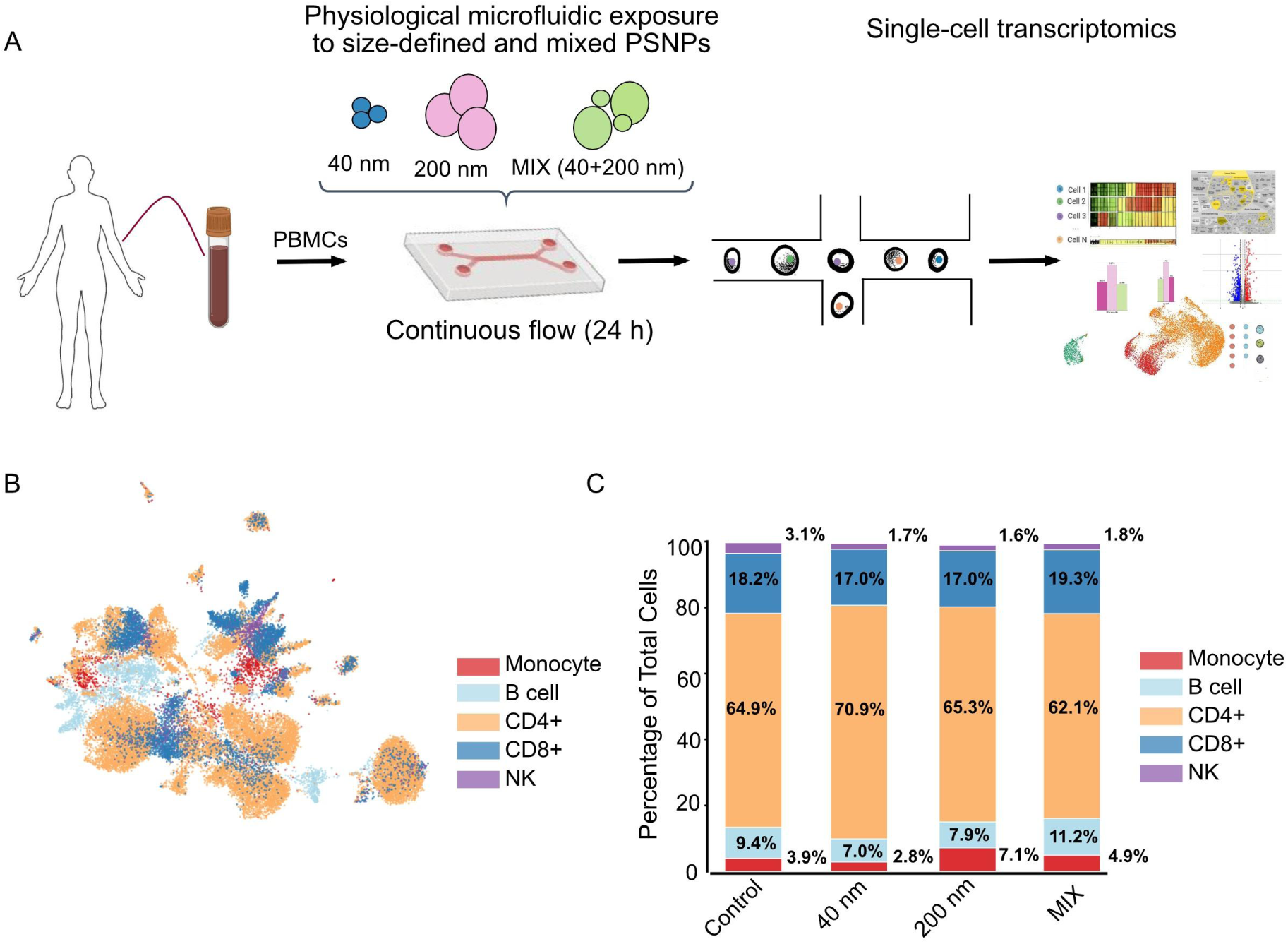
Experimental setup and single-cell landscape of human PBMCs under dynamic exposure to polystyrene nanoparticles (PSNPs). **(A)** Schematic overview of the microfluidic continuous-flow exposure system (24-hour incubation under a constant flow rate with 40 nm, 200 nm, or mixed (40+200 nm) PSNPs). **(B)** Uniform Manifold Approximation and Projection (UMAP) embedding of 33,814 identified cells, color-coded by major immune lineages. **(C)** Relative cellular proportions (%) across control and exposure conditions within the single-cell dataset.

Following exposure, immune cells were subjected to droplet-based scRNA-seq, yielding a high-quality transcriptomic dataset spanning multiple immune cell populations. Reference-based cell type annotation identified major immune cell types, including CD14⁺ monocytes (from hereafter monocytes), B cells, CD4⁺ T cells, CD8⁺ T cells, and a smaller population of natural killer (NK) cells (Figure 1B). Uniform manifold approximation and projection (UMAP) visualization showed clear separation of immune cell identities (Figure 1B), indicating robust annotation across the dataset. To enable unbiased comparison across exposure conditions, single-cell datasets were integrated using Harmony^23^, minimizing condition-driven segregation (Supplementary Figure S1A). All major immune cell populations were recovered across conditions with sufficient cell numbers for cell-type–resolved analyses (Supplementary Figure S1B).

Quantitative analysis of immune cell composition revealed that relative abundances of major cell types were largely preserved across all PSNP exposure conditions (Figure 1C). While minor variations in relative cell-type percentages were observed, these did not follow a systematic pattern with respect to particle size or mixed exposure. These data indicate that PSNP exposure under flow does not disrupt immune cell composition or lineage topology within the examined time frame. Together, these data indicate that nanoplastic exposure does not alter immune composition but instead reprograms transcriptional states in a cell-type-specific manner.

### 3.2 Nanoplastics size differentially modulates transcriptional responses across immune populations

To assess the global transcriptional impact of PSNP exposure, we quantified the number of differentially expressed genes (DEGs) for each immune cell type and exposure condition relative to controls (Figure 2A; Supplementary Table S1). Differential expression analysis was performed using uniform statistical thresholds across all comparisons (adjusted p-value < 0.05).

**Figure 2.**
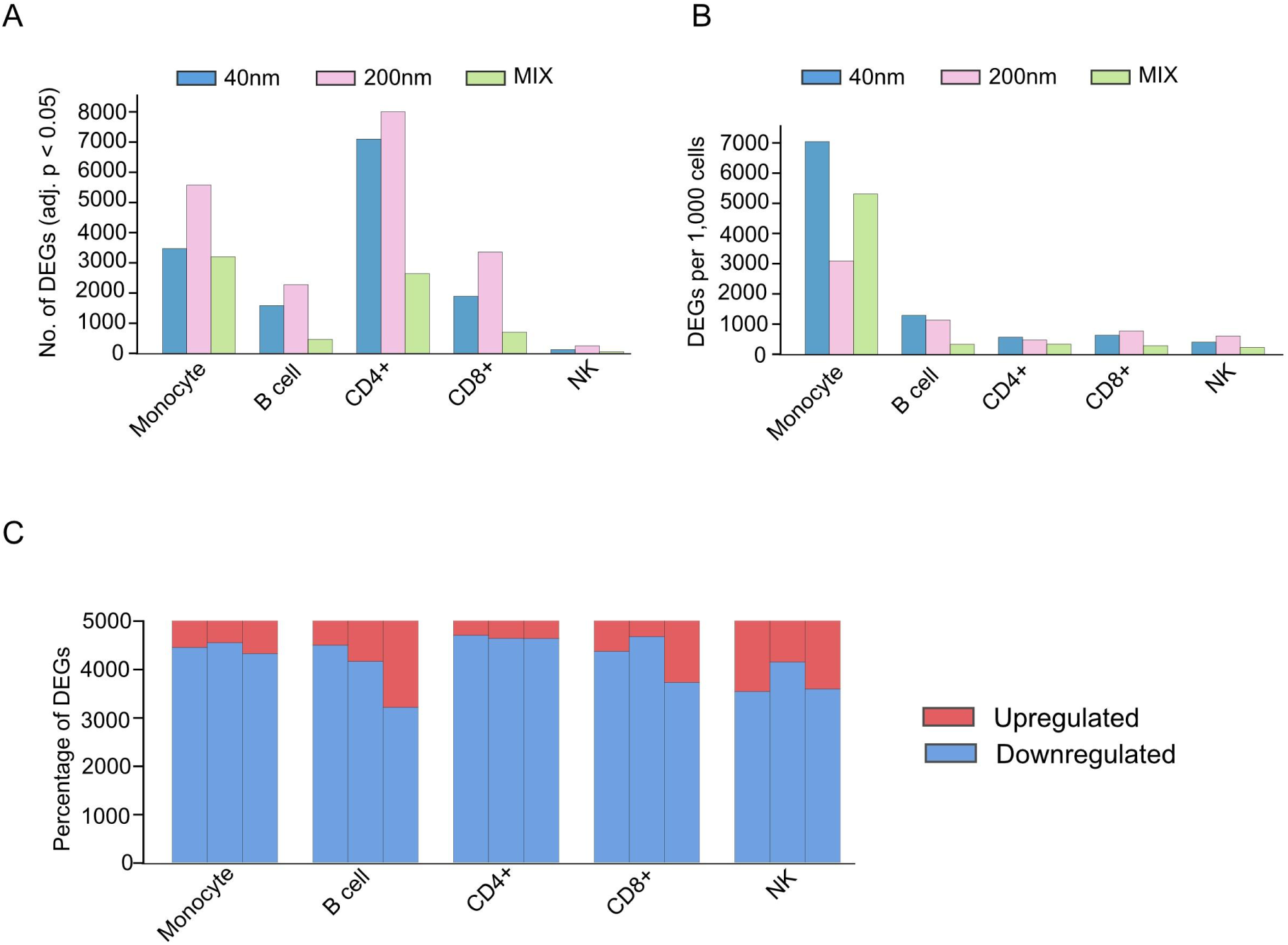
Global transcriptional impact of PSNP exposure across immune cell types. **(A)** Number of DEGs identified in each immune cell type following exposure to 40 nm, 200 nm, or mixed (40 + 200 nm) PSNPs relative to unexposed controls. Statistical significance evaluated using a two-sided Wilcoxon rank-sum test with Benjamini-Hochberg adjustment, adjusted p < 0.05. **(B)** DEG counts normalized to cell abundance, shown as DEGs per 1,000 cells, to account for differences in population size and enable comparison of intrinsic transcriptional responsiveness across immune lineages. **(C)** Directionality of transcriptional changes across immune cell types and exposure conditions, showing the relative proportions of upregulated (red) and downregulated (blue) genes among identified DEGs.

Exposure to 200 nm PSNPs yielded the highest number of DEGs across most immune cell types, exceeding those observed after exposure to 40 nm PSNPs. This trend was evident across both innate and adaptive immune populations, while NK cells showed a comparatively lower DEG burden. Mixed exposure to 40 + 200 nm PSNPs did not yield a simple additive effect; instead, it produced an intermediate or distinct number of DEGs, depending on the cell type. These observations indicate that transcriptional responsiveness varies by particle size and exposure complexity across immune lineages.

Because immune cell populations differ substantially in abundance, we next normalized DEG counts to cell number to calculate DEG density per 1,000 cells (Figure 2B). This analysis revealed that monocytes and NK cells exhibit a disproportionately high transcriptional response relative to their abundance, whereas B cells and T cells display lower DEG density despite contributing more cells to the dataset. Normalization, therefore, reveals intrinsic differences in per-cell transcriptional sensitivity across immune compartments.

Analysis of DEG directionality showed a striking predominance of downregulated genes across immune cell types and exposure conditions (Figure 2C). This bias toward transcriptional repression was consistent across all cell types and particle sizes. In contrast, upregulated genes were relatively few and typically exhibited larger fold changes. Inspection of full volcano plots confirmed this asymmetric response profile across conditions and immune populations (Supplementary Figure S3). Together, these patterns suggest coordinated transcriptional adjustment rather than broad inflammatory activation.

Quality control analyses confirmed that DEG patterns were not driven by systematic differences in sequencing depth, gene detection, or mitochondrial transcript content across conditions (Supplementary Figure S2). Taken together, these analyses demonstrate that PSNP exposure induces size-dependent and exposure-specific transcriptional responses across immune cell types, with innate immune populations exhibiting greater per-cell sensitivity under physiological flow conditions.

### 3.3 Nanoplastic exposure induces a conserved ribosome-associated core program across immune populations

To identify higher-order biological programs underlying the observed transcriptional changes, pathway enrichment analysis was performed across immune cell types and exposure conditions. A curated set of KEGG pathways representing core cellular and immune functions was used to construct a pathway-by-cell-type heatmap (Figure 3A). Across all immune populations and exposure conditions, pathways related to ribosomal function and RNA regulation emerged as a shared signature. This conserved response indicates that modulation of ribosome-associated and RNA-regulatory pathways is a common cellular response to PSNP exposure under flow. The persistence of this signal across innate and adaptive lineages suggests a core intracellular adaptation framework largely independent of particle size.

**Figure 3.**
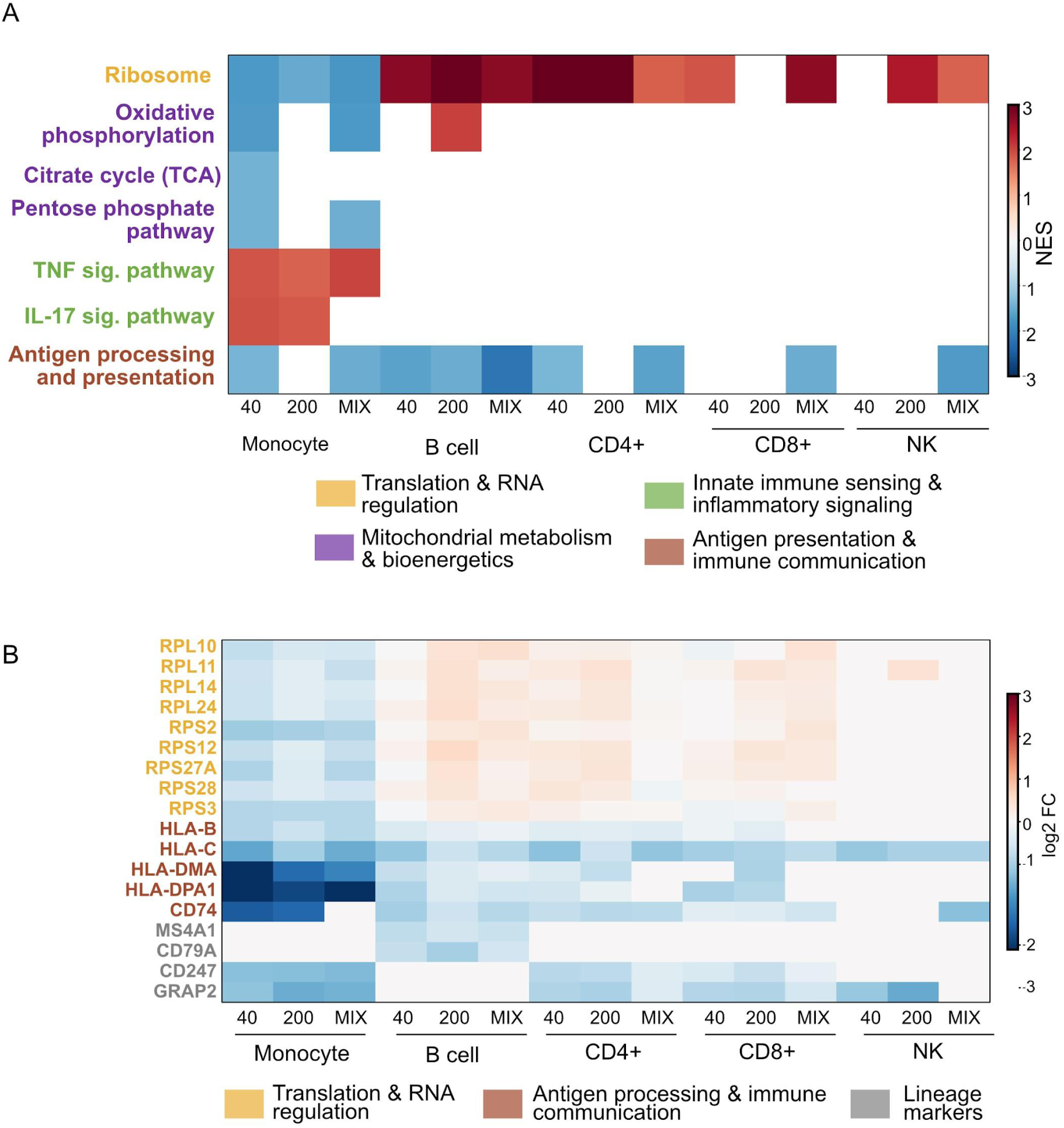
Lineage-specific pathway modulation under single and mixed PSNP exposures. **(A)** Heatmap representing Normalized Enrichment Scores (NES) from GSEA across major immune subsets. Red blocks denote positive pathway enrichment; blue blocks denote negative pathway enrichment (FDR < 0.05). **(B)** Gene-level heatmap displaying relative expression changes (log2 FC) of representative leading-edge genes involved in ribosomal architecture, antigen processing, and lineage markers.

In contrast, pathways related to mitochondrial metabolism and bioenergetics, including oxidative phosphorylation and the citrate cycle, were selectively enriched in monocytes and exhibited clear size dependence (Figure 3A). These pathways were minimally affected in adaptive immune cells, highlighting a divergence between innate and adaptive transcriptional responses. Redox-associated biosynthetic pathways such as the pentose phosphate pathway were likewise enriched predominantly in monocytes, consistent with metabolic rewiring rather than overt inflammatory activation. Inspection of the full KEGG enrichment landscape (Supplementary Table S2) confirms that metabolic and redox-associated programs are primarily engaged in monocytes across exposure conditions (Supplementary Figure S4), supporting cell-type–specific metabolic adaptation.

Canonical inflammatory signaling pathways, including TNF and IL-17 signaling, were selectively enriched in monocytes and were largely absent from adaptive immune cells. Enrichment of these pathways did not coincide with robust induction of classical pro-inflammatory cytokine genes, suggesting regulated signaling engagement rather than acute inflammatory activation. Pathways associated with antigen processing and presentation were detected across multiple immune cell types, reflecting shared immune communication functions while retaining cell-type–specific patterns.

To complement pathway-level analysis, representative genes associated with conserved and cell-type–biased transcriptional programs were examined using a gene-level heatmap (Figure 3B). Genes involved in ribosomal structure and translation (e.g., RPL and RPS family members) showed broad and coherent modulation across immune populations, whereas genes associated with antigen processing and immune communication (e.g., *HLA* genes and *CD74*) were modulated across multiple cell types without forming a dominant shared program. Lineage marker genes were included solely to facilitate interpretation of cell-type–specific expression patterns and were not interpreted functionally.

Together, these analyses demonstrate that PSNP exposure induces a conserved ribosome-associated and RNA-regulatory framework across immune cell types, upon which cell-type–specific metabolic and signaling programs are differentially superimposed. This hierarchical organization underlies the size-dependent transcriptional remodeling observed in monocytes and the broader modulation observed in adaptive immune cells.

### 3.4 Monocytes undergo coordinated metabolic state remodeling in a nanoplastic size-dependent manner

Monocytes displayed the strongest transcriptional sensitivity to PSNP exposure after normalization for cell abundance (Figure 2B), prompting detailed analysis of monocyte-specific transcriptional organization. KEGG pathway enrichment analysis revealed that monocyte responses differed according to PSNP size and exposure complexity (Figure 4A). Exposure to 40 nm PSNPs preferentially enriched pathways associated with mitochondrial metabolism and ribosome-associated regulation, including oxidative phosphorylation, citrate cycle, pentose phosphate pathway, fatty acid elongation, and ribosome-associated programs. In contrast, exposure to 200 nm PSNPs showed stronger enrichment of inflammatory signaling pathways, most prominently TNF signaling, IL-17 signaling, and cytokine–cytokine receptor interaction pathways. Mixed exposure resulted in concurrent enrichment of metabolic and inflammatory programs, indicating distinct combined-exposure transcriptional responses rather than simple amplification of single-particle effects.

**Figure 4.**
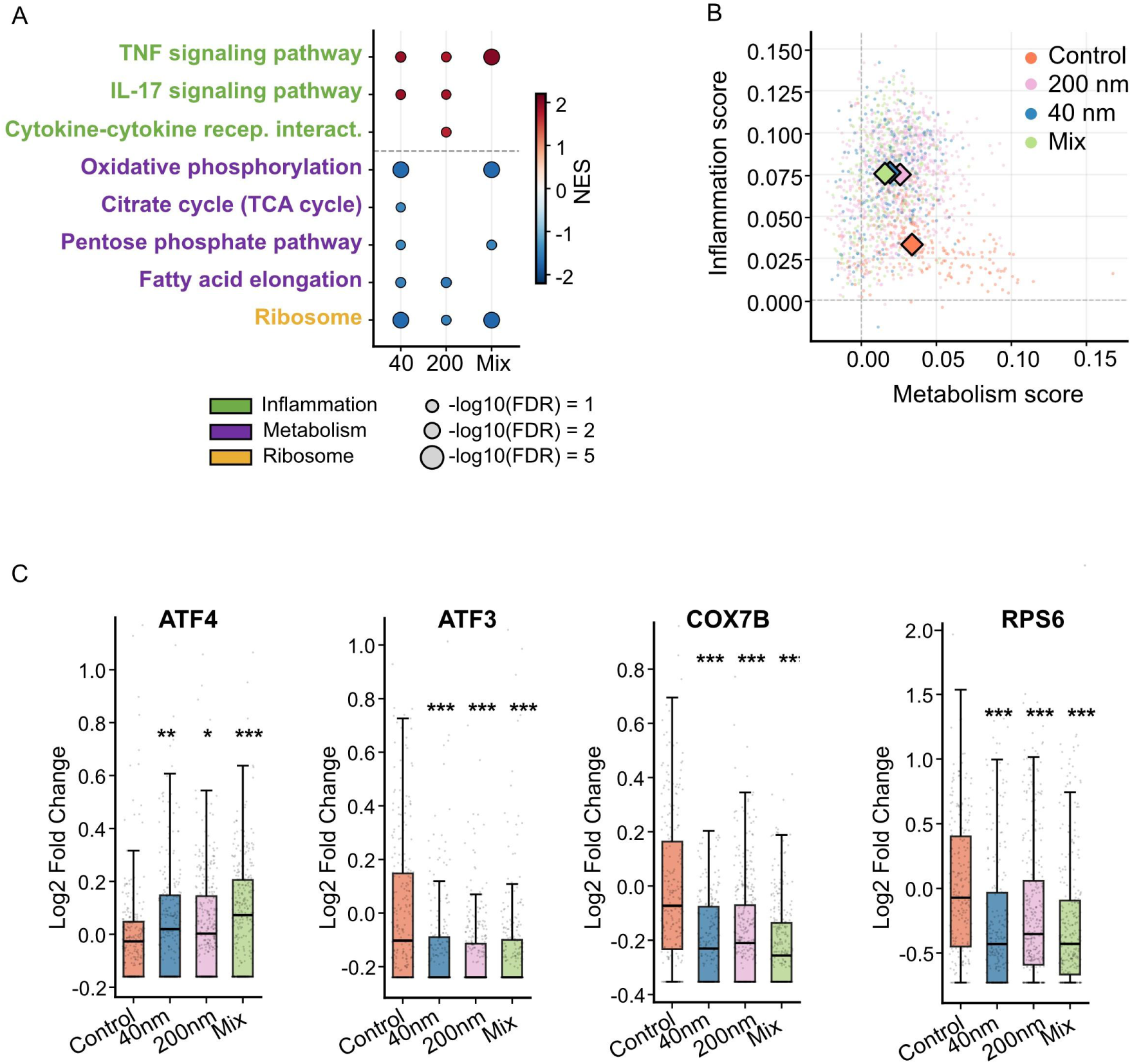
Characterization of the immunometabolic and integrated stress response in monocytes. **(A)** Targeted GSEA profile demonstrating negative enrichment (suppression) of core mitochondrial pathways (oxidative phosphorylation, TCA cycle, pentose phosphate and fatty acid pathways) and ribosome-associated regulation alongside positive enrichment of inflammatory cascades (TNF, IL-17 signaling and cytokine–cytokine receptor interaction pathways). **(B)** Dual-axis scatter plot mapping individual monocyte coordinates based on aggregate metabolic and inflammatory module scores derived via Seurat’s AddModuleScore. Centroids represent condition medians. Metabolic score was calculated from oxidative phosphorylation, TCA cycle, pentose phosphate, and fatty acid pathways, while inflammatory score was calculated from TNF, IL-17, and cytokine–cytokine receptor interaction pathways. **(C)** Expression of representative stress adaptation (*ATF3* and *ATF4*), mitochondrial (*COX7B*), and translational (*RPS6*) remodeling genes in monocytes across exposure conditions. Statistical significance was assessed using Welch’s t-test versus control (* p < 0.05, ** p < 0.01, *** p < 0.001).

To further examine pathway-level organization at single-cell resolution, metabolic and inflammatory module scores were calculated for individual monocytes (Figure 4B). Monocyte populations showed substantial overlap across exposure conditions, with only modest shifts in pathway activity distributions. These observations support coordinated transcriptional modulation in response to PSNP exposure rather than the emergence of discrete inflammatory cell states.

Analysis of the representative DEGs further supported coordinated stress-associated and metabolic transcriptional remodeling in monocytes, including stress-response transcription factors (*ATF3* and *ATF4*), mitochondrial and redox-associated genes (*COX7B, GPX3,* and *SLC25A37*), and translational regulators (*RPS6* and *RPL3*) (Figure 4C and Supplementary Figure S5). In parallel, monocytes displayed selective induction of inflammatory mediators, including IL1B, IL6, PTGS2, and CCL20, together with regulatory genes such as *SIGLEC7* and *FGL2*, indicating controlled inflammatory engagement embedded within broader transcriptional adaptation rather than overt inflammatory activation.

Consistent with preservation of global immune organization, PAGA analysis demonstrated maintenance of inter-lineage connectivity across all exposure conditions without evidence of major topology disruption (Supplementary Figure S6).

Crucially, the co-exposure cohort (Mix) demonstrated a distinct, non-linear threshold effect regarding monocyte energy expenditure. Despite containing half the concentration of the single 40 nm exposure (50 μg/mL vs. 100 μg/mL), the Mix induced a metabolic and translational suppression profile statistically indistinguishable from the pristine 40 nm treatment, as evidenced by highly overlapping negative enrichment scores in oxidative phosphorylation (40 nm NES: −1.71; Mix NES: −1.72) and ribosomal translation (40 nm NES: −1.72; Mix NES: −1.74).

### 3.5 Adaptive immune cells preserve lineage-associated transcriptional organization following PSNP exposure

In contrast to monocytes, adaptive immune cells displayed comparatively modest and lineage-preserving transcriptional responses following PSNP exposure (Figure 5). KEGG pathway enrichment analysis in B cells and CD4+ T cells identified enrichment of ribosome-associated and oxidative phosphorylation pathways across exposure conditions, indicating limited ribosome-associated and metabolic modulation (Figure 5A). In parallel, pathways associated with antigen recognition and immune communication, including B cell receptor signaling, T cell receptor signaling, antigen processing and presentation, cell adhesion molecules, and hematopoietic cell lineage pathways, were reduced across exposure conditions. Analysis of the combined exposure data revealed a non-linear, synergistic suppression of B-cell functional competence. The negative enrichment of both B-cell receptor signaling (BCR) (NES −1.99) and Antigen processing and presentation (NES −2.19) in the Mix dropped significantly below the inhibitory metrics observed in either the 40 nm (BCR: −1.60; Antigen processing and presentation: −1.61) or 200 nm (BCR: −1.51; Antigen processing and presentation: −1.49) exposures.

**Figure 5.**
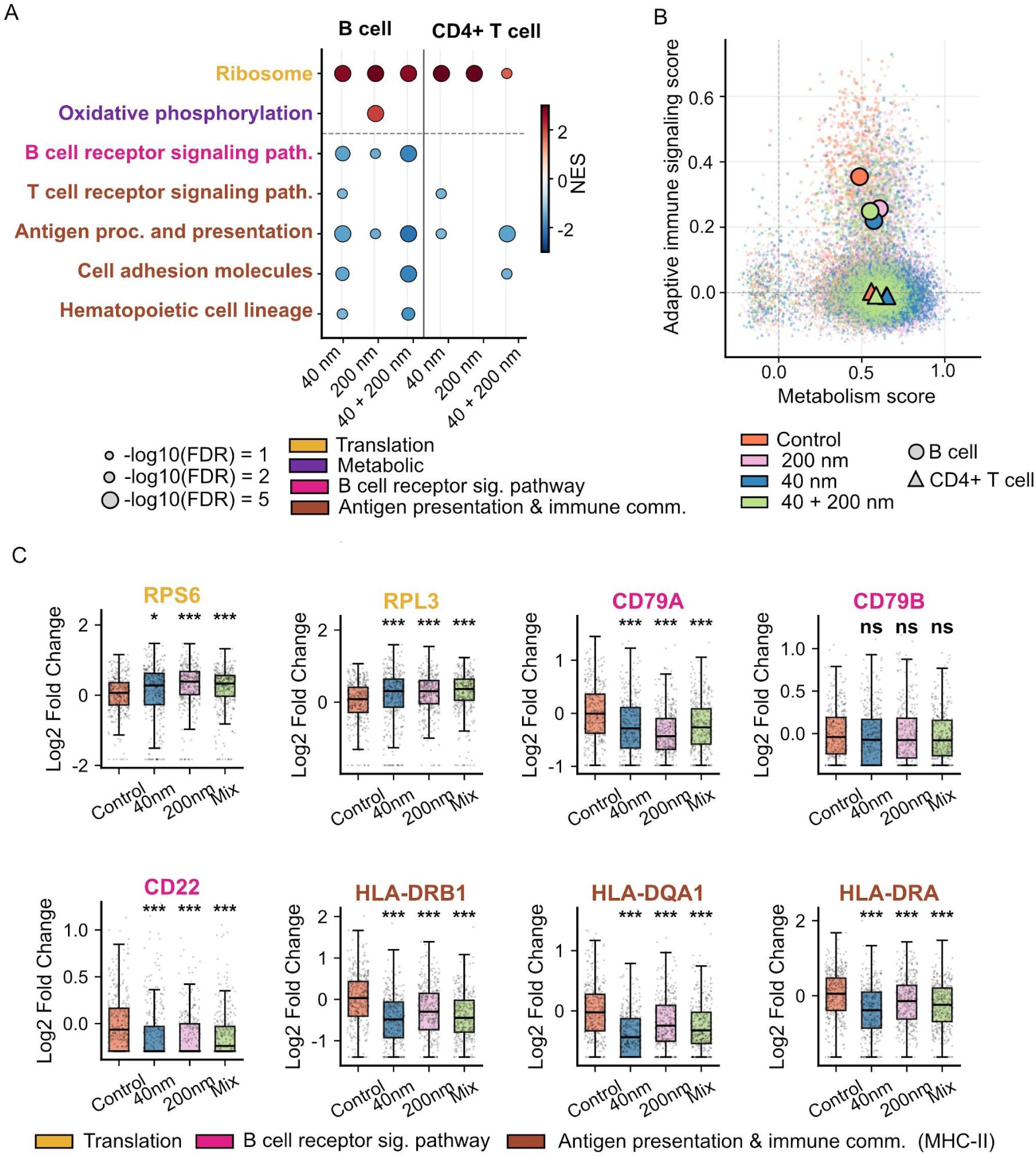
Adaptive immune cells preserve lineage-associated transcriptional organization following PSNP exposure. **(A)** KEGG pathway enrichment analysis of B-cell and CD4+ T-cell transcriptional responses following exposure to 40 nm, 200 nm, or mixed (40 + 200 nm) PSNPs. Ribosome-associated and OXPHOS pathways showed enrichment across exposure conditions, whereas pathways related to antigen recognition and immune communication displayed reduced activity. Dot size represents pathway significance (−log10 FDR), and color indicates NES. **(B)** Per-cell relationship between pathway-derived metabolic capacity and adaptive immune signaling scores in adaptive immune cells. Metabolic capacity was calculated using ribosomal and oxidative phosphorylation signatures, while the adaptive immune signaling score was derived from antigen receptor signaling and MHC-associated pathways. **(C)** Expression of representative ribosomal, antigen presentation, and B-cell receptor–associated genes across exposure conditions. Statistical significance was assessed using Welch’s t-test versus control (* p < 0.05, ** p < 0.01, *** p < 0.001; ns, not significant).

To further examine the relationship between metabolic activity and adaptive immune functionality, module scores were calculated using ribosomal and oxidative phosphorylation signatures (“metabolic capacity”) and antigen receptor/MHC-associated pathways (“adaptive immune signaling score”) (Figure 5B). Adaptive immune populations showed broad overlap across exposure conditions, without evidence of discrete transcriptional state transitions or lineage disruption.

Representative gene expression patterns in B cells further supported preservation of lineage identity despite coordinated transcriptional modulation (Figure 5C). Ribosomal genes, including *RPS6* and *RPL3*, were consistently upregulated across exposure conditions, whereas genes associated with antigen presentation and B-cell lineage-associated identity, *including HLA-DRA, HLA-DQA1, HLA-DRB1, CD22,* and *CD79A*, showed reduced expression. Together, these findings indicate that adaptive immune cells respond to acute PSNP exposure through modest transcriptional adjustment while preserving overall lineage-associated transcriptional architecture.

### 3.6 Distinct organizational principles characterize innate and adaptive immune responses to PSNP exposure

To integrate the transcriptomic responses observed across diverse circulating blood lineages, we mapped the pathway-level architecture of innate and adaptive immune cohorts following PSNP exposure under dynamic flow (Figure 6). The resulting structural model highlights a compartmentalized divergence in cellular behavior. Monocytes exhibit an uncoupled, intracellular immunometabolic remodeling signature, characterized by the concurrent downregulation of translation (ribosome-associated programs) and bioenergetic cascades alongside the activation of targeted inflammatory pathways. These innate perturbations fluctuate strictly as a function of particle size and exposure complexity. Conversely, adaptive lymphocytes (CD4⁺ T and B cells) preserve their basic metabolic parameters and structural lineage identity, displaying a localized, superficial suppression of antigen-receptor signaling and intercellular communication pathways. Together, these data indicate that innate and adaptive immune compartments organize transcriptomic responses through fundamentally distinct mechanisms under physiological circulation conditions.

**Figure 6.**
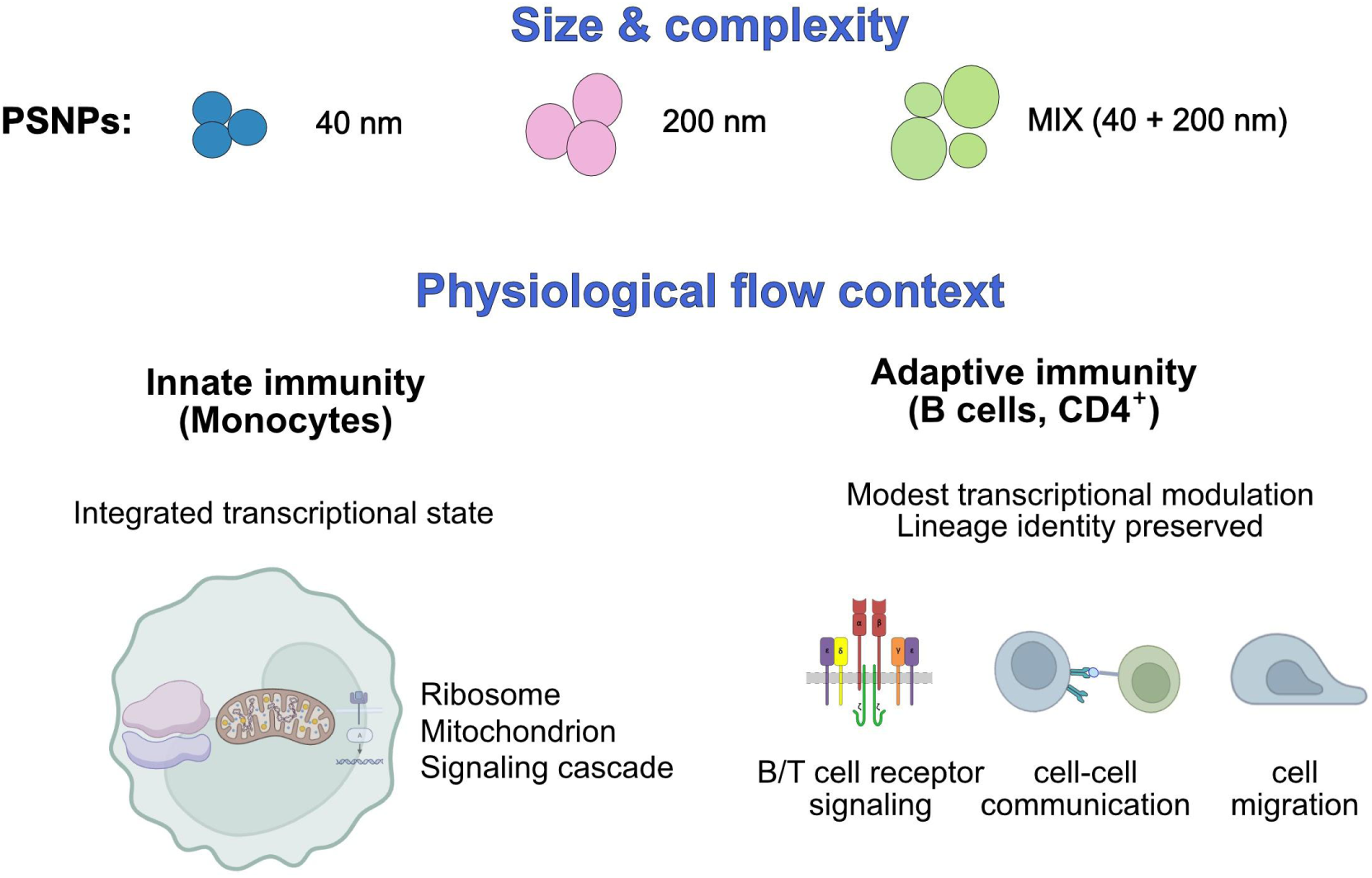
Distinct transcriptional organization patterns characterize innate and adaptive immune responses to PSNP exposure. Schematic representation of transcriptional response patterns observed across immune compartments following exposure to PSNPs of different sizes (40 nm, 200 nm) and their combination (40 + 200 nm) under physiological flow conditions. Monocytes displayed coordinated modulation of ribosome-associated, mitochondrial, and inflammatory signaling pathways, which varied with particle size and exposure complexity. In contrast, adaptive immune cells exhibited comparatively modest transcriptional modulation involving antigen receptor signaling, immune communication, and cell interaction pathways while preserving lineage-associated transcriptional organization.

## 4. DISCUSSION

This study combines a continuous-flow microfluidic exposure platform with single-cell transcriptomics to investigate how nanoplastic size and exposure complexity shape the transcriptome landscape of human PBMCs. A key finding is that acute PSNP exposure under physiological flow conditions does not significantly change overall cellular composition, but instead induces compartment-specific transcriptomic remodeling rather than a uniform, pan-leukocyte immune response. Monocytes emerged as the most sensitive population, displaying highly coordinated, pathway-level changes when accounting for cell abundance. This highlights that absolute DEG counts alone do not adequately capture the higher-order biological organization under particulate stress.

Distinct transcriptomic profiles emerged as a function of particle size and mixture complexity. Exposure to monodisperse 40 nm PSNPs was characterized by a negative enrichment of core mitochondrial bioenergetic and biosynthetic cascades, including oxidative phosphorylation and the citrate cycle. Crucially, co-exposure to 40 + 200 nm PSNPs (Mix) demonstrated a non-linear threshold effect on monocyte energy expenditure. Despite containing half the concentration of the single 40 nm exposure, the Mix induced metabolic and translational (ribosome) suppression nearly identical to that of the pristine 40 nm treatment. This indicates that 40 nm-containing exposures showed the most pronounced metabolic signatures, in which low relative doses of the smaller fraction are sufficient to trigger a metabolic downregulation.

Conversely, pristine 200 nm PSNPs preferentially engaged downstream immune signaling pathways, including the TNF and IL-17 cascades. Under mixed exposure conditions, the concurrent engagement of these independent metabolic and inflammatory programs suggests a unique transcriptional architecture of combined exposure. This multi-size co-existence is highly relevant to realistic exposure scenarios, where immune cells encounter heterogeneous particulate populations rather than monodisperse materials, indicating that immune responses to complex nanoplastic environments are not fully predictable from simplified single-particle models. Because the mixed exposure matrix inherently halved the effective mass concentration of each individual particle size relative to the monodisperse treatments (50 μg/mL vs. 100 μg/mL), future investigations incorporating systematic, multi-tier dose-response matrices are required to completely isolate concentration-dependent thresholds from responses driven by physical, heterogeneous particle-particle integration.

This uncoupled immunometabolic phenotype in monocytes, characterized by active inflammatory transcripts alongside a suppressed bioenergetic engine, can be molecularly explained by a targeted uncoupling within the Integrated Stress Response (ISR) architecture. Across all exposure profiles, monocytes exhibited a distinct uncoupled signature marked by the upregulation of the emergency stress-sensor transcript *ATF4* alongside the concomitant downregulation of its core homeostatic feedback repressor, *ATF3*. This ‘broken brake’ configuration could explain why the inflammatory response remained comparatively constrained at the global transcriptional level during the examined exposure window, supporting a temporal model in which intra-organelle transcriptional reorganization precedes overt systemic immune activation.

The observed coupling between metabolic containment, ribosome-associated downregulation, and inflammatory pathway organization in monocytes structurally resembles features of trained immunity. In classical trained immunity, innate cells undergo long-term functional reprogramming, characterized by altered metabolic activity and context-dependent inflammatory responsiveness (Ochando et al., 2023). Although the present study does not address long-term innate memory or secondary challenges, the acute coupling observed here suggests that PSNP exposure may engage early adaptive features of innate immune reprogramming under physiological flow conditions.

In contrast to the intense intracellular remodeling of monocytes, adaptive immune cells exhibited a comparatively modest, lineage-preserving transcriptional modulation. While B cells and CD4⁺ T cells demonstrated differential gene expression, these changes remained dispersed across superficial immune communication, receptor signaling, and migratory pathways while preserving fundamental inter-lineage organization. Nonetheless, localized pathway analysis of the combined exposure data revealed a highly non-linear, synergistic suppression of B-cell functional competence. The downregulation of both the B-cell receptor signaling cascade and the antigen processing and presentation pathway in the Mix cohort significantly exceeded the inhibitory metrics observed in either single-size treatment alone. This suggests that complex, multi-size particulate systems reinforce the operational blockade of the adaptive apparatus beyond a simple additive effect, effectively arresting the cellular machinery needed for antigen recognition and lymphocyte coordination.

The coordinated transcriptional adjustments observed under dynamic flow complement and extend recent proteomic and cellular evidence detailing how nanoplastic size shapes myeloid polarization and oxidative stress. Previous studies using static exposure systems demonstrated that PSNPs preferentially affect monocyte-lineage cells and induce oxidative stress, mitochondrial dysfunction, and inflammatory signaling. Wolff et al. reported that monocyte-derived macrophages and dendritic cells were substantially more sensitive to PS particles than adaptive immune populations (Wolff et al., 2023), whereas Fusco et al. demonstrated a preferential association of PSNPs with monocytes and dendritic cells, accompanied by oxidative and mitochondrial stress responses (Fusco et al., 2025). The present study, by examining primary human PBMCs under controlled physiological flow at single-cell resolution, uncovers the upstream, pathway-level architecture that underpins these downstream phenotypes. Rather than inducing uncoordinated, overt cytotoxic stress, dynamic conditions reveal a highly organized immunometabolic containment, characterized by coordinated stress-response and inflammatory pathway engagement.

The induction of *ATF4*-associated stress programs, together with modulation of oxidative phosphorylation, the pentose phosphate pathway, and the citrate cycle, further supports convergence with previously described nanoplastic-induced metabolic stress responses. This emphasizes the validity of mechanistic investigation, even though it was conducted using single-donor PBMCs, which was chosen to minimize confounding genetic variance. Together, these findings support a coordinated framework for innate immune transcriptional adaptation during acute PSNP exposure under physiological flow conditions and suggest that dynamic exposure systems may reveal more restrained and organized response states than conventional static culture models.

The deployment of a microfluidic exposure platform represents a critical experimental feature of this work. By maintaining primary human cells under continuous flow, the system constrains material–cell interactions within a dynamic environment that closely approximates physiological circulation, minimizing localized, artificial sedimentation and high-dose artifacts that are dominant in static culture systems. Consequently, the dynamic exposure conditions optimized here reveal more restrained, coordinated pathway-level adjustments. This distinction is particularly vital for nanoscale materials, whose behavior in static systems fails to recapitulate the kinetic interactions that occur in intravascular circulation.

While this investigation focused on polystyrene nanoplastics, the size-dependent transcriptional organization patterns identified here are likely applicable to other nanoscale materials with varying dimensions and surface interaction properties. These data establish physiological flow as a critical parameter in modeling material–immune interactions, identify monocyte metabolic programming as a primary vulnerability axis, and provide a conceptual framework for understanding how nanoscale material parameters shape systemic immune coordination within dynamic physiological environments.

## 5. Environmental implication

Environmental exposure to micro- and nanoplastics occurs as complex mixtures of particles differing in size, concentration, and physicochemical properties. While most current toxicological studies rely on static exposure systems and bulk analyses, the present work demonstrates that nanoplastic exposure under physiological flow conditions induces a coordinated, cell-type–specific transcriptional organization in human immune cells. In particular, monocytes displayed integrated modulation of ribosome-associated, metabolic, and inflammatory signaling pathways without evidence of initiating immediate cytolytic or decompensated hyper-inflammatory states, suggesting that acute nanoplastic exposure may induce an early homeostatic reorganization of innate immune function prior to the development of classical toxicological phenotypes. Although environmental exposure is generally considered chronic, acute high-particle-load exposure scenarios generated by consumer products and food-contact materials remain poorly understood, particularly at the level of immune transcriptional organization under physiological flow conditions. The observation that combined exposure to different particle sizes engages concurrent transcriptional programs further highlights the importance of considering heterogeneous environmental nanoplastic mixtures rather than simplified single-particle models. These findings support integrating physiologically relevant flow systems and single-cell approaches into future environmental nanotoxicology assessment frameworks.

A limitation of the present study is that all experiments were performed using PBMCs from a single healthy donor. Although the observed preferential responsiveness of monocytes is consistent with previous reports of enhanced nanoplastic interactions with monocyte-lineage cells, inter-individual variability could influence the magnitude and organization of transcriptional responses. Therefore, the present findings should be considered a high-resolution characterization of nanoplastic-induced immune adaptation under physiological flow conditions and warrant validation across larger donor cohorts and additional exposure paradigms.

## 6. CONCLUSION

This study demonstrates that the immunotoxic effects of PSNPs on human PBMCs are characterized by lineage-specific transcriptional reprogramming rather than a uniform cellular response. By combining microfluidic continuous-flow systems with high-resolution single-cell transcriptomics, we resolve distinct mechanical vulnerabilities across circulating immune subsets under physiologically relevant dynamic conditions.

The primary conclusions of this work are summarized below:

- **Compartment-Specific Vulnerability:** Innate and adaptive immune compartments exhibit fundamentally divergent responses to nanoparticle stress. Monocytes emerge as the primary target of acute toxicological injury, undergoing a profound coordinated downregulation of core mitochondrial bioenergetic and translation machineries. In contrast, adaptive lymphoid lineages (CD4⁺ T and B cells) maintain their metabolic baseline and lineage identity but become functionally inactivated.
- **The Immunometabolic Paradox:** Monocytes display a decoupled transcriptional signature under PSNP stress, marked by the concurrent activation of inflammatory signaling networks alongside a repressed metabolic baseline. At the single-cell level, this uncoupled phenotype is defined by the upward mobilization of the emergency stress-sensor *ATF4*, accompanied by the suppression of its homeostatic feedback regulator *ATF3*, demonstrating that acute exposure induces an unbraked state of transcriptional cell exhaustion.
- **Non-Linear Mixture Dynamics:** Co-exposure to mixed-size nanoparticle populations triggers cell-type–specific non-linearities that cannot be predicted using conventional monodisperse linear modeling. Within the innate compartment, 40 nm particles exhibit clear size-dependent dominance, with lower comparative fractions sufficient to reach a toxicological threshold and cause metabolic shutdown. Concurrently, within the adaptive compartment, multisize interactions drive highly synergistic, compounding suppression of antigen presentation networks and B-cell receptor signaling.
- **Methodological Relevancy:** The integration of continuous microfluidic flow minimizes the localized, artificial sedimentation artifacts common to static culture systems. The resulting data reveal a more controlled, coordinated, and pathway-level immunometabolic adaptation, indicating that dynamic exposure environments provide a more precise representation of early circulating blood-cell interactions.

Collectively, these findings underscore that evaluating nanoplastic hazards purely through mass-based or single-size linear baselines systematically underestimates the biological complexity of composite particulate stress. High-resolution single-cell profiling of heterogeneous, multi-size matrices is critical to accurately mapping the true systemic risks posed by emerging plastic contaminants to human health.

## CRediT authorship contribution statement

V. K. Software, Formal analysis, Visualization, Writing: original draft.

M. M. Conceptualization, Formal analysis, Writing: original draft, Writing: review and editing.

N. M. D. Conceptualization, Methodology, Investigation, Funding acquisition, Data curation, Formal analysis.

M. Z. Formal analysis.

M. I. Formal analysis.

Z. Bioinformatics analysis and visualization.

M. G. J. Formal analysis.

K. Formal analysis.

U. P. Z. Supervision. F.P. Supervision.

N. F. Funding acquisition, Project administration.

Lj. Conceptualization, Methodology, Investigation, Data curation, Funding acquisition, Project administration, Supervision. All authors: Writing, review, and editing.

## Declaration of generative AI and AI-assisted technologies in the manuscript preparation process

During the preparation of this work, the authors used ChatGPT (OpenAI) to edit language and refine the manuscript text. After using this tool/service, the author(s) reviewed and edited the content as needed and take full responsibility for the content of the published article. Claude code Opus 4.7 was used as an assistant for generating visualizations.

