## Supplementary figures for "Heterogeneous nanoplastic exposure induces integrated immunometabolic states in human monocytes under physiological flow"

[1]School of Electrical Engineering, University of Belgrade, Bulevar Kralja Aleksandra 73, Belgrade, Serbia;
[2]Institute for Information Technologies, University of Kragujevac, Jovana Cvijića bb, 34000, Kragujevac, Serbia;
[3]Institute of Molecular Genetics and Genetic Engineering (IMGGE), University of Belgrade, Vojvode Stepe 444a, 11042 Belgrade, Serbia;
[4]Faculty of Science, University of Kragujevac, Radoja Domanovića 12, Kragujevac;
AZ:
[5]Faculty of Medical Sciences, Department of Genetics, University of Kragujevac, Svetozara Markovića 69,34000 Kragujevac, Serbia;
[6]Faculty of Pharmacy, University of Belgrade, Vojvode Stepe 450, 11221 Belgrade, Serbia;
[7]BIA Separations CRO, Labena, Teslova 30, 1000 Ljubljana, Slovenia,
UPZ:
[8]Labena d.o.o., Belgrade, Serbia,
FP:
[9]Faculty of Engineering, University of Kragujevac, Sestre Janjic 6, 34000 Kragujevac, Serbia;

[+]These authors contributed equally to this work

**Keywords:** polystyrene nanoplastics; microfluidics; scRNA-seq; PBMCs; monocytes; immunometabolism

### Supplementary Figures

Fig. S1

A

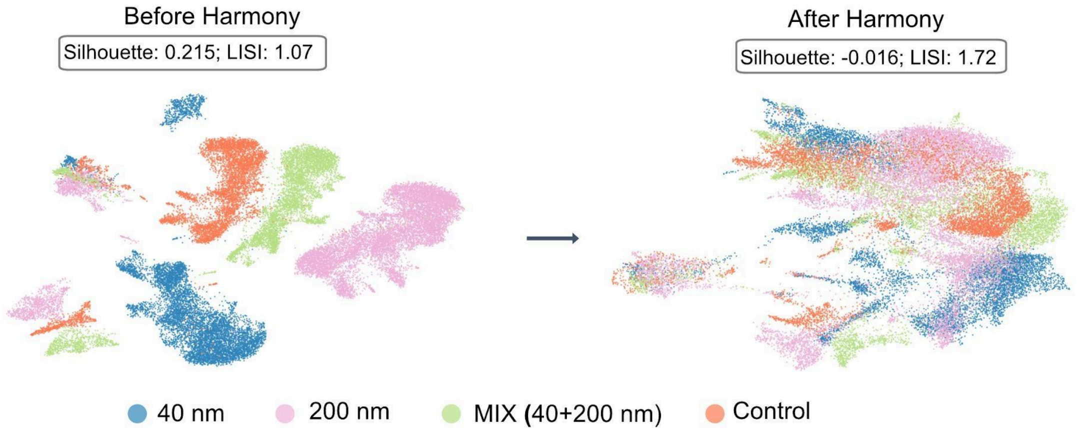


B

| | Control | 40 nm | 200 nm | MIX |
|---|---|---|---|---|
| Monocytes | 257 | 246 | 900 | 301 |
| B cells | 615 | 611 | 999 | 688 |
| CD4+ | 4229 | 6186 | 8272 | 3826 |
| CD8+ | 1185 | 1483 | 2157 | 1190 |
| NK | 205 | 149 | 205 | 110 |

**Supplementary Figure S1. Computational batch correction, multi-condition data integration, and lineage-specific cell recovery. (A)** Uniform Manifold Approximation and Projection (UMAP) embedding of 33,814 human peripheral blood mononuclear cells (PBMCs) before (left) and after (right) dataset integration utilizing the Harmony algorithm. Cell coordinates are color-coded by experimental exposure condition (Control, 40 nm, 200 nm, and mixed (40 + 200 nm) PSNPs). Internal tracking metrics denote the absolute Global Silhouette scores and Local Inverse Simpson's Index (LISI) calculations across both integration processing states. **(B)** Matrix table enumerating the exact number of quality-controlled single cells successfully recovered and analyzed per distinct immune lineage, stratified by the four separate experimental exposure conditions.

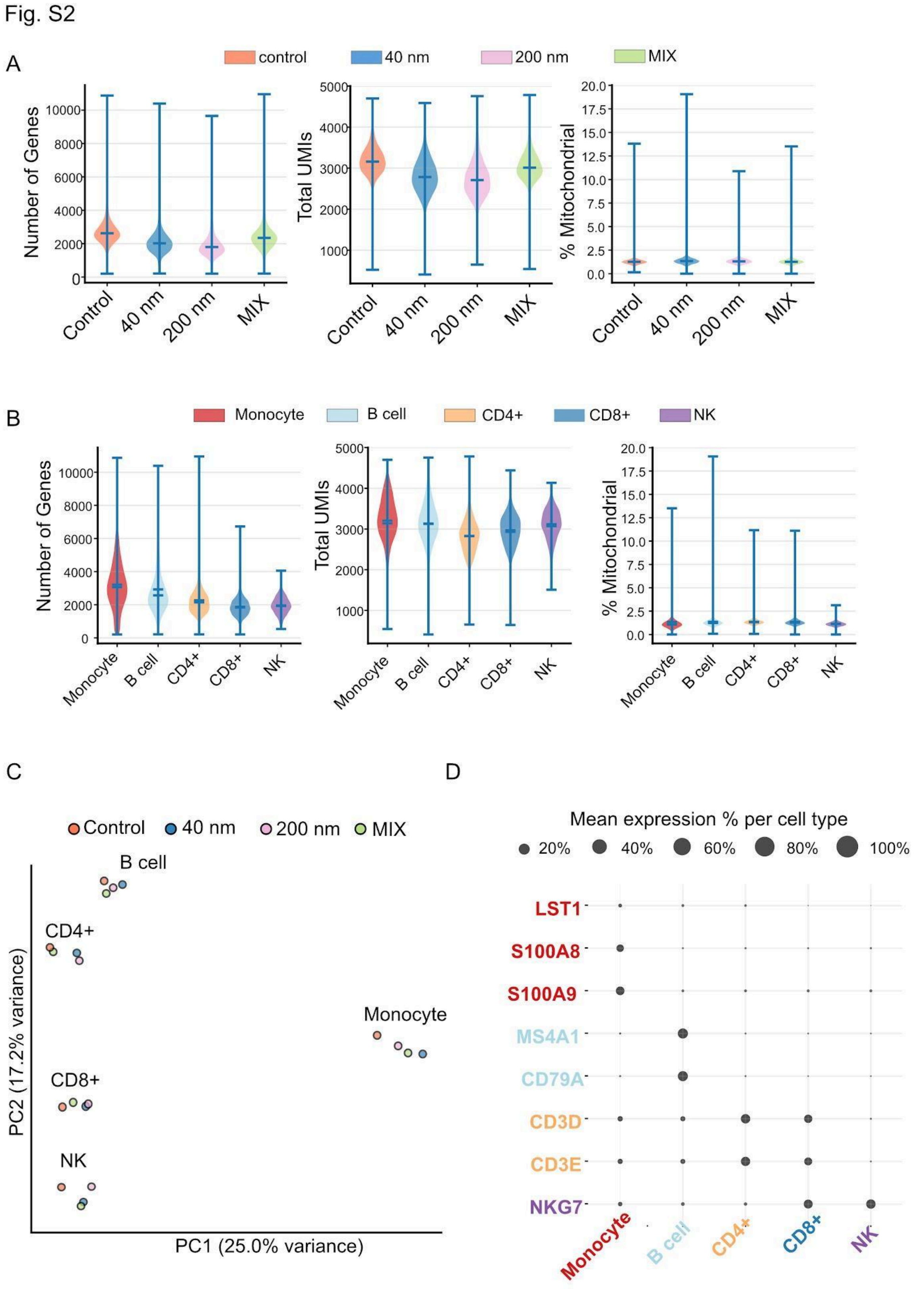


**Supplementary Figure S2. Single-cell RNA sequencing quality control filters, principal component separation, and cluster marker validation. (A)** Single-cell quality control distribution metrics across individual exposure conditions (Control, 40 nm, 200 nm, Mix), evaluating the absolute number of detected genes per cell, total unique molecular identifier (UMI) counts, and the exact percentage (%) of transcripts mapping to the mitochondrial genome. **(B)** Quality control distribution profiles stratified natively across annotated cell lineages (Monocyte, B cell, CD4$^{+}$ T cell, CD8$^{+}$ T cell, NK cell). **(C)** Principal Component Analysis (PCA) plot mapping the transcriptomic variance and global separation of the five discrete, annotated immune populations. **(D)** Dot plot validating cluster annotations using canonical transcript markers (**LST1, S100A8, S100A9, MS4A1, CD79A, CD3D, CD3E, NKG7**). Dot diameter indicates the percentage of expressing cells, and color intensity denotes relative mean log-normalized expression values.

Fig.S3

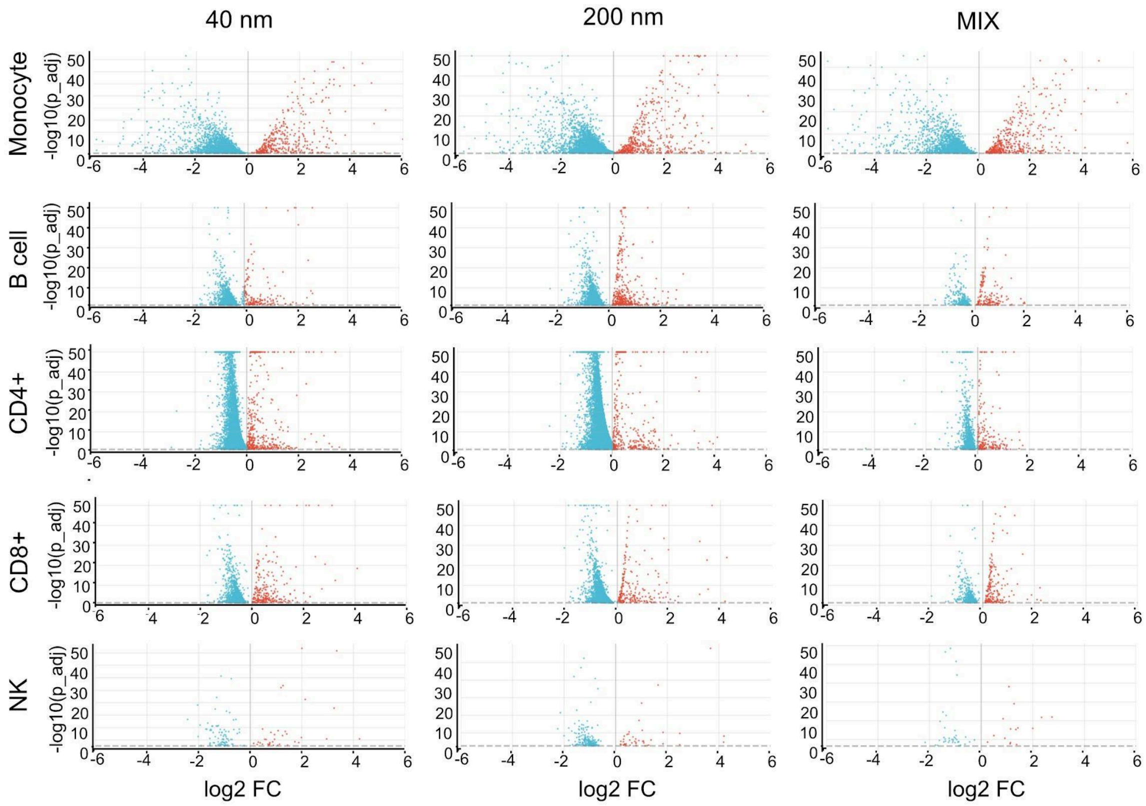


**Supplementary Figure S3. Differential gene expression landscapes across primary immune lineages following monodisperse and mixed PSNP exposure.** High-resolution volcano plots displaying global differentially expressed gene (DEG) profiles for major immune cell populations (monocytes, B cells, CD4$^{+}$ T cells, CD8$^{+}$ T cells, and NK cells) following acute exposure to 40 nm, 200 nm, or mixed (40 + 200 nm) PSNPs compared directly to unexposed control cells. The horizontal axis plots the absolute log2 fold change (log2 FC), and the vertical axis plots the associated status of statutory significance -log 10 adjusted p-value (-log10 (p_adj). Red dots denote significantly upregulated transcripts; blue dots denote significantly downregulated transcripts based on uniform, false discovery rate (FDR)-adjusted thresholds.

Fig. S4

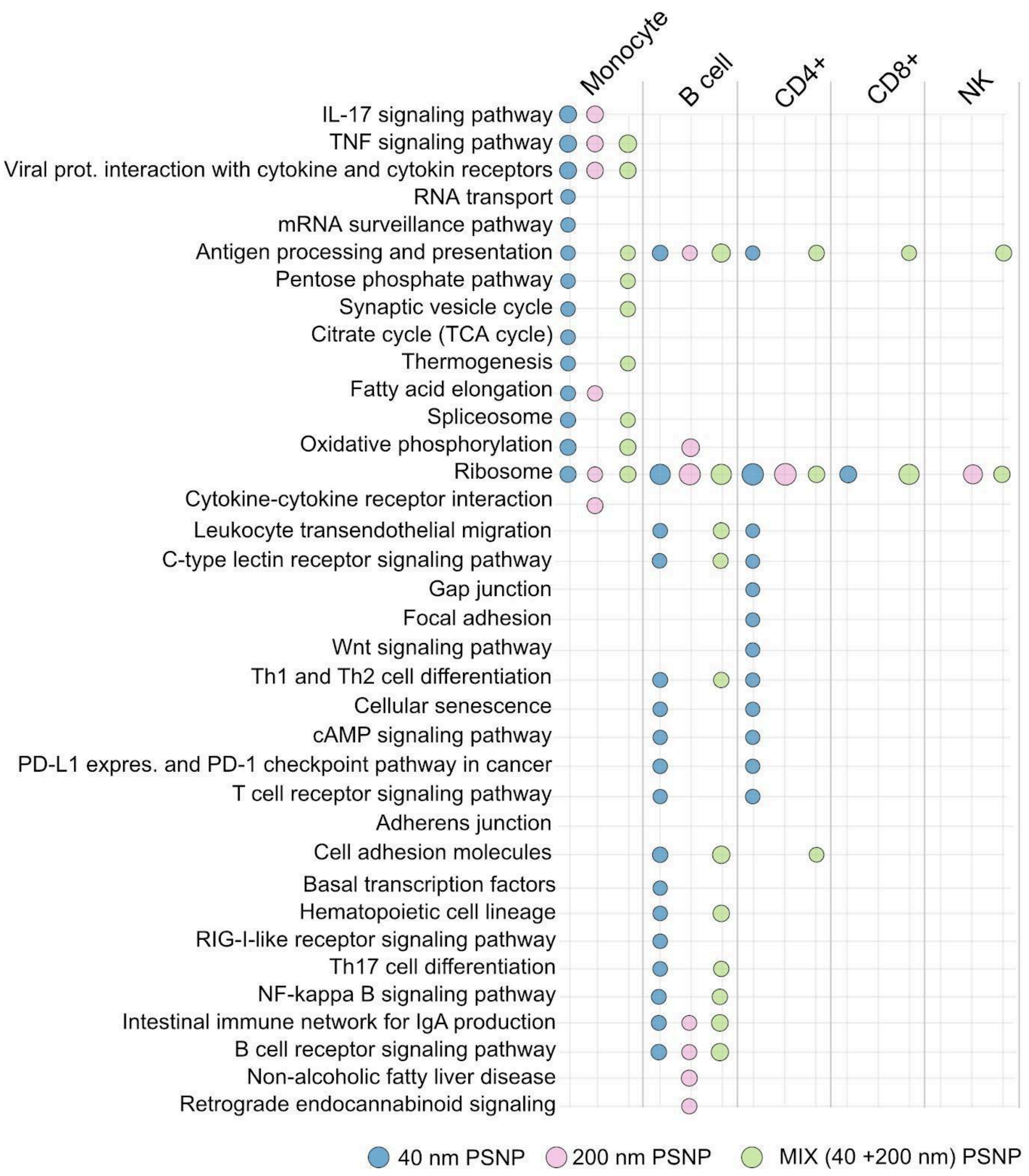


**Supplementary Figure S4. Comprehensive uncollapsed KEGG pathway enrichment profiling across circulating leukocyte lineages.** Dot plot summarizing the complete, uncollapsed Kyoto Encyclopedia of Genes and Genomes (KEGG) pathway enrichment landscape resolved across major immune cell subsets and individual PSNP exposure profiles. Circular plot markers are color-coded strictly by specific treatment conditions: blue denotes 40 nm PSNP, pink denotes 200 nm PSNP, and green denotes mixed (40 + 200 nm) PSNP treatments. Visual plotting presence indicates statistical significance based on an adjusted threshold of (FDR < 0.05)

Fig. S5

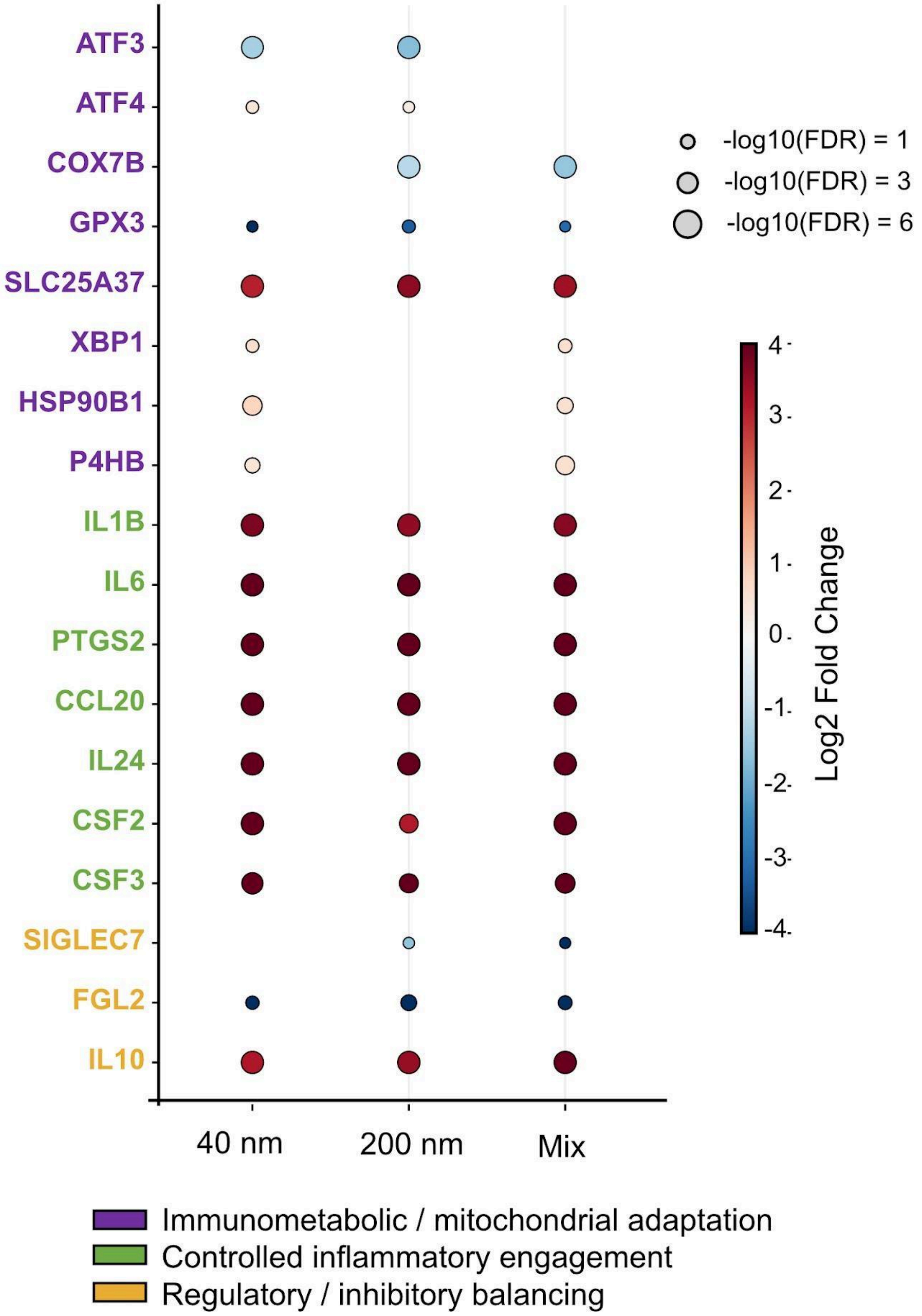


**Supplementary Figure S5. Target gene expression profiles highlighting coordinate immunometabolic, inflammatory, and regulatory shifts in monocytes.** Dot plot visualizing specific log-normalized fold transformations (log2 FC) of core leading-edge transcripts across the 40 nm, 200 nm, and mixed exposure models in the monocyte compartment. Gene clusters are structurally segregated by functional modules: Immunometabolic/mitochondrial adaptation, Controlled inflammatory engagement, and Regulatory/inhibitory balancing. Circle circumferences represent specific false discovery rate (FDR) significance thresholds, and the continuous color gradient maps the precise directional magnitude of the expression changes.

Fig. S6

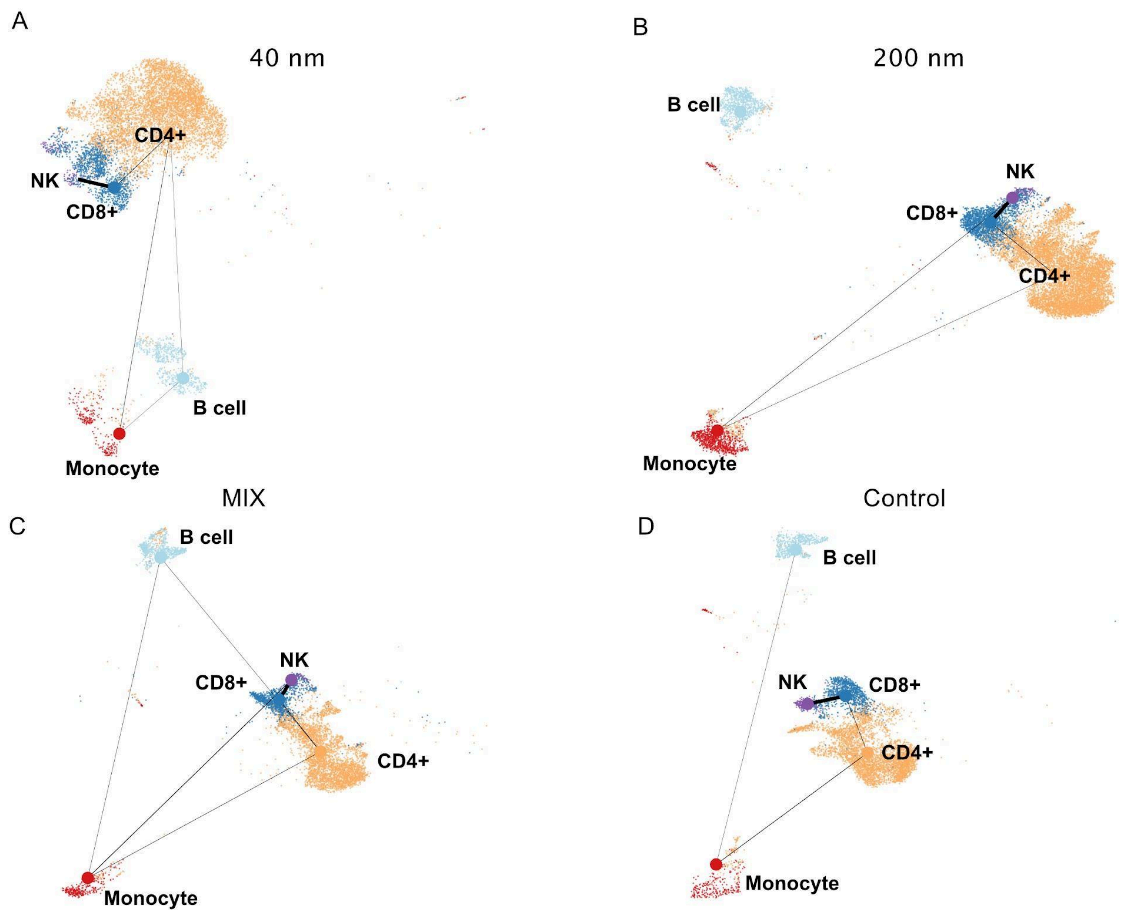


**Supplementary Figure S6. Partition-based graph abstraction (PAGA) modeling of immune lineage topology and connectivity under nanoparticle stress.** Network topology graphs derived from PAGA depicting the mathematically inferred structural connectivity and lineage relationships among major immune lineages under distinct experimental conditions: **(A)** 40 nm PSNP, **(B)** 200 nm PSNP, **(C)** Mixed (40 + 200 nm) PSNP, and **(D)** unexposed control. Circular nodes represent distinct immune cell clusters, and the thickness of the interconnecting vector edges indicates the absolute calculated structural connectivity strength between major cell lineages.
